# Knock-knock, who’s there: Sex-coding, competition and the energetics of tapping communication in the tok-tok beetle, *Psammodes striatus* (Coleoptera: Tenebrionidae)

**DOI:** 10.1101/509257

**Authors:** John R.B. Lighton

## Abstract

I describe the abdomino-substratal tapping communication system of a Southern African tenebrionid beetle, *Psammodes striatus* (Fabricius, 1775) (Coleoptera: Tenebrionidae: Molurini), using computer simulation of tapping signals and computer-assisted acquisition of precise response timing data, augmented with data from natural beetle-beetle communication. Communication consists of trains of 5 - 7 Hz taps in groups or trains separated by 2-3 sec intervals. Male beetles spontaneously produce groups of tap-trains with 8 - 18 taps per train. If other beetles reply, an alternating duet commences. Solitary female beetles do not tap spontaneously but respond to male tapping with short, distinctive tap-trains containing 4 – 6 taps; they ignore female signals. In contrast, extensive communication occurs between male beetles, the nature of which changes significantly if the stimulus call is typical of male or of female beetles. Inter-male communication consists of long tap-trains, but males interacting with females produce shorter tap-trains and engage in phonotactic behavior that is absent in inter-male communication. Females respond highly preferentially to inter-male communication, rather than to the signals produced spontaneously by single males. Finally, I propose a simple model of the selective advantages of this unusual communication system, and calculate its approximate energetic leverage over random locomotion (∼13x).

## INTRODUCTION

Communication via substrate vibrations is widespread among insects and is regarded primarily as a mechanism for communication between potential mates (Greenfield 2002, Cocroft and Rodriguez 2005; see also reviews by Hill 2001, 2008). Among the coleoptera, vibrational communication or “tapping”, also known as “drumming”, has been described in at least 24 species (summarized by Hill 2008).

Beetle taxa differ in the pulse repetition frequency of tapping, and in male vs. female participation in tapping behavior. In the death watch beetle *Xestobium rufovillosum*, both males and females tap, though males cannot be distinguished from females in terms of tapping characteristics (Birch and Keenlyside 1991; White et al. 1993). Tschinkel and Doyen (1976) found that male tapping behavior in the beetle *Eusattus reticulatus* increases in the presence of females, but that only males tap, and males do not respond to the presence of other males. Similarly, Pearson and Allen (1996) reported that females of another *Eusattus* species also do not tap, and Slobodchikoff and Spangler (1979) found that only males of *Eupsophulus castaneus* tap. However, in the African tenebrionid beetle, *Phryanocolus somalicus*, both males and females tap (Zachariassen 1977; Kristensen and Zachariassen 1980), but, as with the death watch beetle, “the beetles cannot use the sound signals for determination of the sex of other beetles”.

I have quantified the energy cost of tapping communication in the South African molurine tenebrionid *Psammodes striatus*, also known colloquially as the “toktokkie” or tok-tok beetle (Lighton 1987). Using the energy cost data, together with data from an earlier study in which I measured the cost of pedestrian locomotion in this species (Lighton 1985), I calculated that substrate-borne vibrational communication was approximately tenfold more energetically efficient for mate-location than a random walk (Lighton 1987). However, the description of the tapping communication in that paper was limited to a broad description of the tap-trains produced by male beetles only. The purpose of this paper is to refine that description, adding data on male vs. female tapping behavior, and on male-male tapping interactions. A simple model is proposed to explain the selective benefit of male-male communication in this species, and the energy allocation advantages of tapping communication over mate-location by locomotion are explored.

## METHODS AND MATERIALS

### Animals

Males and females of *Psammodes striatus* were collected during October in the austral spring from Sandy Bay (Cape Peninsula, South Africa). All beetles were labeled on one elytron with a small circle of white correcting fluid (Tipp-Ex; Tipp-Ex Company, Frankfurt, Germany) on which a code number was written. The beetles were kept on fine sand and small stones in glass vivaria in an air-conditioned laboratory at the University of Cape Town (22 ± 1 °C, with a natural day/night lighting cycle) and were separated by sex. Oats and lettuce were supplied ad libitum. Beetles maintained in this way remained in apparent good health and were de-labeled and released at the site of capture after the study was concluded.

### Tap-monitoring and analysis techniques

A brief description of tapping behavior is necessary to place the monitoring methodology in context. Close observation in the field and the laboratory established that the beetles tapped in “trains” of ∼4-20 taps, depending on sex and circumstance. Tap-trains were separated by a short pause. Furthermore, tap-trains occurred in “groups”. Each group consisted of a variable number of tap-trains, separated in time from each other by no more than 5-10 seconds, as opposed to the much longer and more variable pauses between groups of trains. The beetles communicate with each other by alternating tap-trains (beetle A – beetle B – beetle A, etc.) in duets. If no inter-beetle communication is taking place, male beetles will occasionally produce spontaneous groups of tap-trains.

The train of taps was chosen as the basic unit for monitoring purposes. Because inter-tap pauses within tap-trains were nearly constant (see Results), a train of taps could be accurately described by its duration and the number of taps comprising it. Further, by measuring the time elapsed between trains, each train could be placed in temporal relation with its neighbors.

The beetle being monitored was kept in a glass-walled terrarium (300 x 220 x 220 mm i.d.) with a 50 mm deep layer of fine sand covering its base. An electret microphone was buried in the top layer of this sand. A 300 x 220 mm sheet of 3mm thick medium density fiberboard (Masonite), on which the beetle had complete freedom of movement, was placed on the sand in contact with the body of the microphone. The terrarium was placed on a heavy granite-topped balance table to isolate it from acoustic interference. The output of the microphone was fed into a variable-gain amplifier, then into a Schmitt trigger and an adjustable monostable multivibrator. The gain of the amplifier was adjusted so that the weakest tap produced by the beetle under study would trip the Schmitt trigger. This sensitivity did not change significantly over the entire surface of the board. The monostable was designed to give a 10 msec output pulse, preventing the counting of multiple pulses from each impact. The resulting pulse was fed into an Acorn BBC computer running monitoring software written by the author.

If a tap was detected that was not followed by a further tap within 400 ms (about twice the normal inter-tap period within a tap-train) it was ignored. Such single signals were always caused by electrical transients or mechanical disturbances. If the tap formed part of a train, the time of the first tap was stored, and further taps were counted until 400 ms had elapsed without the detection of a further tap. The time from the first tap (less 400 ms), and the time elapsed from the end of the penultimate tap-train to the first tap of the last train were stored in the computer, together with the number of taps in the last train. All times were measured with a resolution of 10 msec. These data were regularly transferred to disc together with date, time and observational notes, allowing a complete reconstruction of tapping activity during the monitoring period.

Stimulation was applied to the beetles as computer-synthesized “taps” which consisted of 4 msec bursts of 666 Hz squarewaves. These were fed via a 0.5W power amplifier into a 50 mm permanent-magnet speaker placed cone down on the Masonite floor of the monitoring container. The amplifier gain was adjusted so that a soft but audible tap was produced. The number of taps produced together with their period, length and spacing were controlled by the computer. An inter-tap period of 170 ms was adopted throughout the experimentation, as this was close to the mean inter-tap period of both male and female beetles, and initial experiments showed negligible response effects with moderate changes in tap period. Marked changes in response occurred when the number of taps per train was altered, so this parameter was chosen as the sole variable in the presented stimulus for this study.

Observation of interacting beetles revealed a stereotyped alternation of response between tappers (see Results). A ‘conversation’ was initiated by unstimulated tapping by one beetle, to which another beetle replied with a train of taps. The first beetle would then reply with a second train of taps, and so on. The computer mimicked this behavior. Initially, it synthesized a single train of taps every minute. If the beetle responded, the stimulus was re-presented 1.50 s after the completion of the beetle’s “reply”. This value was chosen as a compromise between the typical delays shown by male and female beetles (see Results), to ensure inter-comparability of results between runs. The beetle then replied again, followed by the computer, and so on. As such ‘conversations’ could continue almost indefinitely (>3 h), a standardized monitoring format was adopted in which the beetle was first stimulated for 20 minutes (the “stimulated state”), following which stimulation stopped (the “poststimulated state”). The poststimulus period ended ten minutes after stimulation stopped, or - if the beetle continued producing trains of taps after this period - until at least 5 seconds had elapsed between trains. The computer could produce a fixed number of stimulus taps per train, or change the number of stimulus taps at random within specified limits. Stimulus taps per train were varied at random between 2 and 18 inclusive (with a constant number of taps within each monitoring period) for data discussed in this paper.

### Statistics

All means are accompanied by their standard deviations and N. The significance criterion employed for all tests was p < 0.05. All statistical software was written by the author and validated against example data sets from Sokal and Rohlf (1973) and Bailey (1959). For comparisons between small (n < 100) data sets with similar variances, Student’s *t* test was used. Variances were tested for significant difference with the F test. Where variances differed significantly but sample sizes were below 100, the approximate t test was used (Bailey 1959). Frequency distributions of data are displayed as probability distributions rather than standard histograms. “Probability” is used in the sense of the maximum likelihood estimator (i.e. P[n] = [number of occurrences of n]/[total number of occurrences]). The area under the distribution is equal to unity. This method eliminates differences in sample size. Linear regression was performed by the least-squares method. Where noted in the text, slopes were linearized by appropriate axis transformations. As an index of variability, the coefficient of variation (CV; standard deviation / mean expressed as a percentage) was used.

## RESULTS

The communication behavior of *Psammodes striatus* is highly stereotyped. The beetle, whether male or female, taps or drums on the substrate by elevating its body and then directing it downwards so that the abdominal tergites forcibly contact the substrate. Male beetles, but not females, possess a small patch of plumose setae at the point of contact, the function of which is unknown. Initial exploration of the system uncovered little variation between individuals except on the basis of body mass, which significantly altered tapping frequency (author’s unpublished data). Thus, the beetles used in this study were selected from a larger sample of beetles on the basis of near-identical body masses. This reduced the feasible sample size to three male and two female beetles, mean masses 2.7 ± 0.1 and 3.0 ± 0.1 g respectively. To compare the stimulated communication results to actual male-female interactions, these data were supplemented by two additional pairs of freely communicating male and female beetles in the same body mass range.

### Unstimulated tapping behavior

See Table 1 for a summary of the data for male beetles. Fig. 1 shows the probability distribution of taps per train for female and male beetles.

**Table 1.** Data on *unstimulated* tapping in 3 male beetles (2 monitoring periods of 45 min each per beetle, 18h45 to 17h30 on consecutive evenings). Female beetles did not tap without stimulation.

|  |  |  |  |  |  |  |
| --- | --- | --- | --- | --- | --- | --- |
| 485 | UNIT | MEAN | S.D. | CV | n | TOTAL |
| 486 | taps per minute | 7.77 | 6.57 | 84.6 | 6 | 2898 taps |
| 487 | trains per group | 4.74 | 3.83 | 80.8 | 46 | 218 trains |
| 488 | taps per train | 9.63 | 2.34 | 24.3 | 218 | 1098 taps |
| 489 | tap frequency ( Hz) | 5.54 | 1.40 | 25.6 | 1880 | 2898 taps |
| 490 | inter-train pause (s) | 2.11 | 0.55 | 20.3 | 168 | 168 periods |

**Figure 1.**
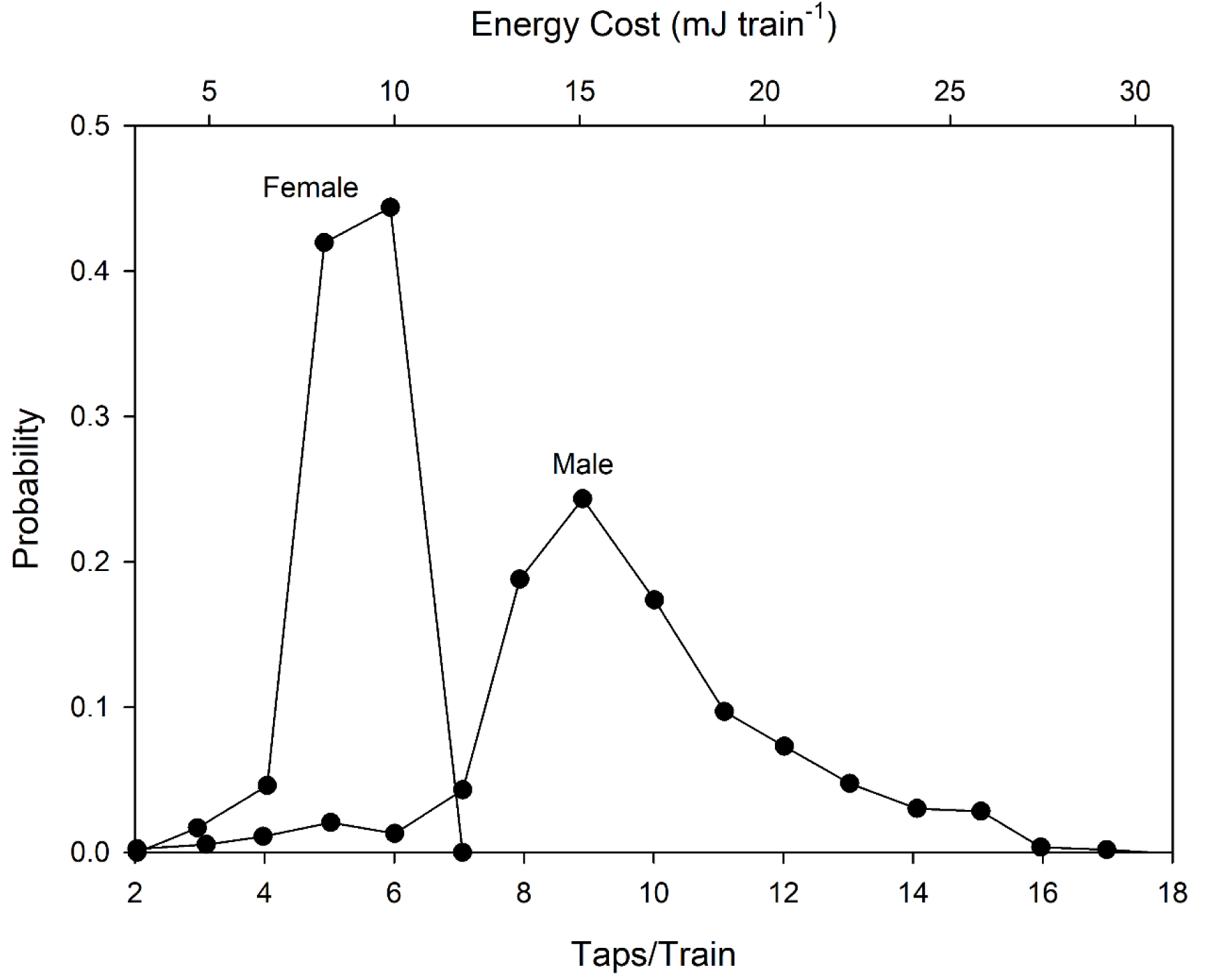
Probability distribution of taps per train in female and male *Psammodes striatus* beetles. n = 2098 taps, 218 trains. See text for definition of these terms. Male tapping was spontaneous (unstimulated). Female tapping was in response to male tapping signals (stimulated), as they did not tap without stimulation. Energy cost in this and subsequent figures is based on 1.725 mJ tap^-1^ for a beetle weighing ∼3 grams (Lighton 1987).

Female beetles did not engage in unstimulated tapping. Tapping by female beetles was fairly rare even when introduced into vivaria with males and constantly stimulated by the males’ calls. After insemination, females responded less often, if at all, to males’ calls (author’s unpublished data). Most duetting interactions were between males, rather than between males and females. No inter-female communication was ever observed even when several females were kept together for long periods.

### Stimulated tapping behavior

A total of 105572 stimulated taps, comprising 4352 trains and 546 groups, delivered by three male and two female beetles during 48 monitoring periods of 30 minutes each, were analyzed. Each beetle was allocated 9-10 monitoring periods. Results are shown in Tables 2 and 3.

**Table 2.** Large-timescale tapping phenomena: a comparison of stimulated and post-stimulated male and female beetles (n = 3 and 2 beetles over 30 and 18 monitoring periods, respectively. SM = Stimulated males; PM = Post-stimulated males; SF = Stimulated females; PF = Post-stimulated females. See text for more details regarding these categories. In two sessions, the females did not respond to the stimulus; these are not included in the analysis.

| 495 | UNIT | MEAN | S.D. | CV | n | TOTAL | BEETLES |
| --- | --- | --- | --- | --- | --- | --- | --- |
| 496 | taps per minute | 100.15 | 25.85 | 25.8 | 30 | 60078 taps | SM |
| 497 | taps per minute | 31.55 | 12.80 | 40.6 | 18 | 11365 taps | SF |
| 498 | taps per minute | 113.82 | 71.74 | 63.0 | 30 | 34129 taps | PM |
| 499 | taps per minute | 2.73 | 2.20 | 80.6 | 18 | 491 taps | PF |
| 500 | groups per minute | 0.442 | 0.176 | 39.8 | 30 | 265 groups | SM |
| 501 | groups per minute | 0.342 | 0.160 | 46.7 | 18 | 123 groups | SF |
| 502 | groups per minute | 0.527 | 0.348 | 66.0 | 30 | 158 groups | PM |
| 503 | trains per group | 16.74 | 11.69 | 69.8 | 263 | 4438 trains | SM |
| 504 | trains per group | 17.22 | 13.62 | 79.1 | 123 | 2106 trains | SF |
| 505 | trains per group | 17.77 | 23.80 | 133.9 | 158 | 2808 trains | PM |

**Table 3.** Short-timescale tapping phenomena. Note the sharply reduced CVs compared to Table 2. The heading S? refers to the grouping applied to the taps. (S>9) refers to 10 or more taps per stimulus train; S<8 to 7 or less; S to the complete range (2-18). ‘Splitting’ by stimulus tap-number was performed on a subset of the data, as explained in the text.

|  |  |  |  |  |  |  |  |  |
| --- | --- | --- | --- | --- | --- | --- | --- | --- |
| 513 | UNIT | MEAN | S.D. | CV | n | TOTAL | S? | BEETLES |
| 514 | tap frequency (Hz) | 5.47 | 1.07 | 19.6 | 55640 | 60078 taps | S | SM |
| 515 | tap frequency (Hz) | 7.42 | 0.47 | 6.4 | 9259 | 11365 taps | S | SF |
| 516 | tap frequency (Hz) | 5.45 | 1.10 | 20.2 | 31321 | 34129 taps | S | PM |
| 517 | taps per train | 15.22 | 2.71 | 17.8 | 1765 | 26864 taps | S>9 | SM |
| 518 | taps per train | 12.24 | 3.40 | 27.8 | 1718 | 21035 taps | S<8 | SM |
| 519 | taps per train | 5.34 | 0.91 | 16.4 | 2106 | 11365 taps | S | SF |
| 520 | taps per train | 12.72 | 2.3 | 18.1 | 1016 | 12928 taps | S>4 | PM |
| 521 | taps per train | 11.14 | 2.21 | 19.8 | 773 | 8611 taps | S<8 | PM |
| 522 | inter-train pause (s) | 2.09 | 0.52 | 24.9 | 1488 | 1488 pauses | S>9 | SM |
| 523 | inter-train pause (s) | 2.19 | 0.67 | 30.6 | 1413 | 1413 pauses | S<8 | SM |
| 524 | inter-train pause (s) | 1.03 | 0.20 | 19.4 | 1482 | 1482 pauses | S | SF |
| 525 | inter-train pause (s) | 3.13 | 0.59 | 18.8 | 829 | 829 pauses | S>9 | PM |
| 526 | inter-train pause (s) | 3.05 | 0.58 | 19.0 | 671 | 671 pauses | S<8 | PM |

Stimulated females tapped with a very narrow frequency distribution (Fig. 1 “Female”, Table 3). The number of response taps per train did not vary as a function of the number of stimulus taps per train (linear regression, *P* > 0.3). The female tap-train appears to be a binary or yes/no response; either given or withheld, depending on the acceptability of the stimulus. Other female call parameters were, however, affected, as shown later.

Males, in contrast to females, showed a wider taps/train distribution and a significantly higher number of taps/train, with the male mode approximately two-fold higher than the female mode (Fig. 1 “Male”, Table 3). The wider distribution is explained by a significant increase in the number of reply taps/train in response to increasing stimulus taps/train (Fig. 2).

**Figure 2.**
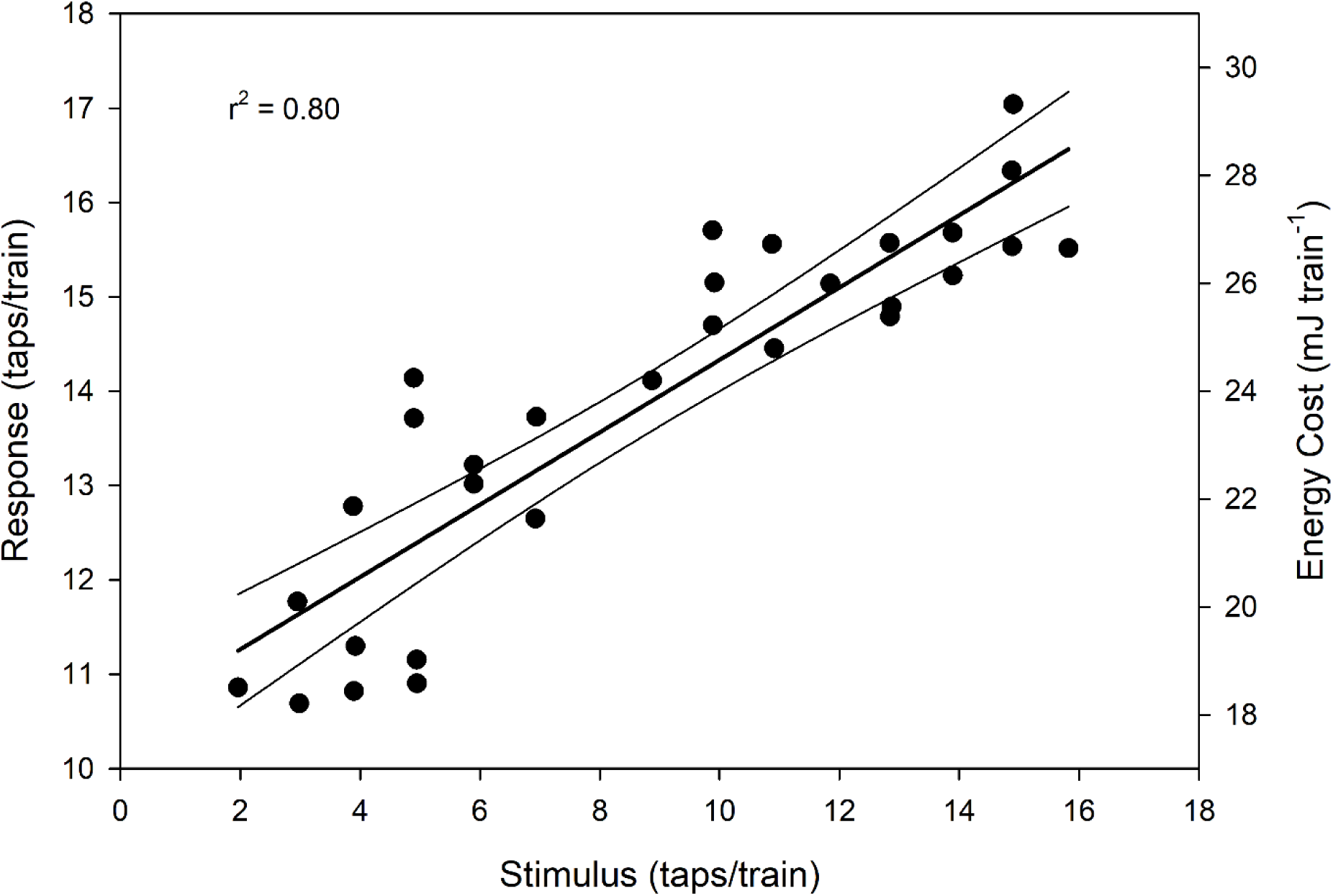
Relation between stimulus tap-number and male response tap-number while stimulus is applied. Female tap-number was monotypic and did not change with stimulus tap number. n = 30 sessions, r^2^ = 0.80, F_1,28_ = 112.63, *P* < 10^-6^; Y = 10.49+0.3835*X.

Males were, however, significantly more likely to engage in protracted duets (number of response tap-trains) if stimulated by tap/train numbers characteristic of female beetles (Fig. 3). As energy expenditure increases linearly with increasing numbers of taps (assuming a constant tap impact and inter-tap period, which is valid; Lighton 1987), this graph also acts as an indication of how much energy is expended by the male beetle as a function of stimulus tap-number.

**Figure 3.**
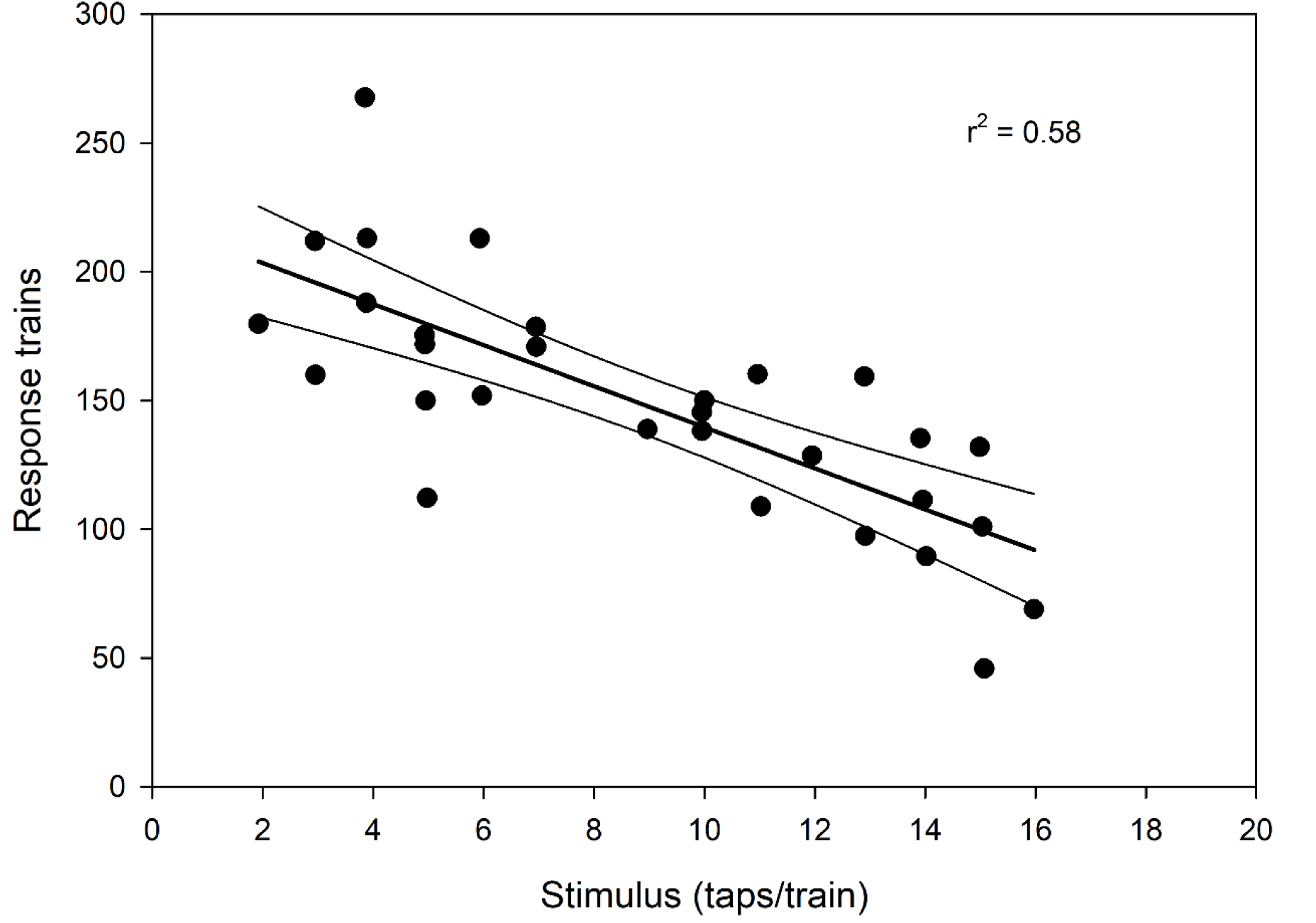
Relation between stimulus tap-number and number of male response trains per monitoring period. n = 30 sessions, r^2^ = 0.58, F_1,28_ = 39.36, *P* < 10^-6^; Y = 219.3-7.972*X.

The males thus showed a distinct response to stimulus tap-trains containing fewer than 8 taps. In addition to tapping persistently in response to such female-characteristic tap-trains, they engaged in phonotactic behavior, which consisted of regular rotational re-orientations of their principal body axis combined with intermittent locomotion, probably serving to alter the amplitude (for linear changes in position) or phase relationships (for rotational changes) of the stimulus signal in a manner informative to a male beetle attempting to locate the source of the stimulus. These behaviors did not occur if the stimulus taps exceeded 8 taps per train. The induction of phonotactic behavior is shown by the sharply increased CV (28 *vs.* 18%; Table 3) in taps/train when responding to signals characteristic of female beetles; see also Fig. 4.

**Figure 4.**
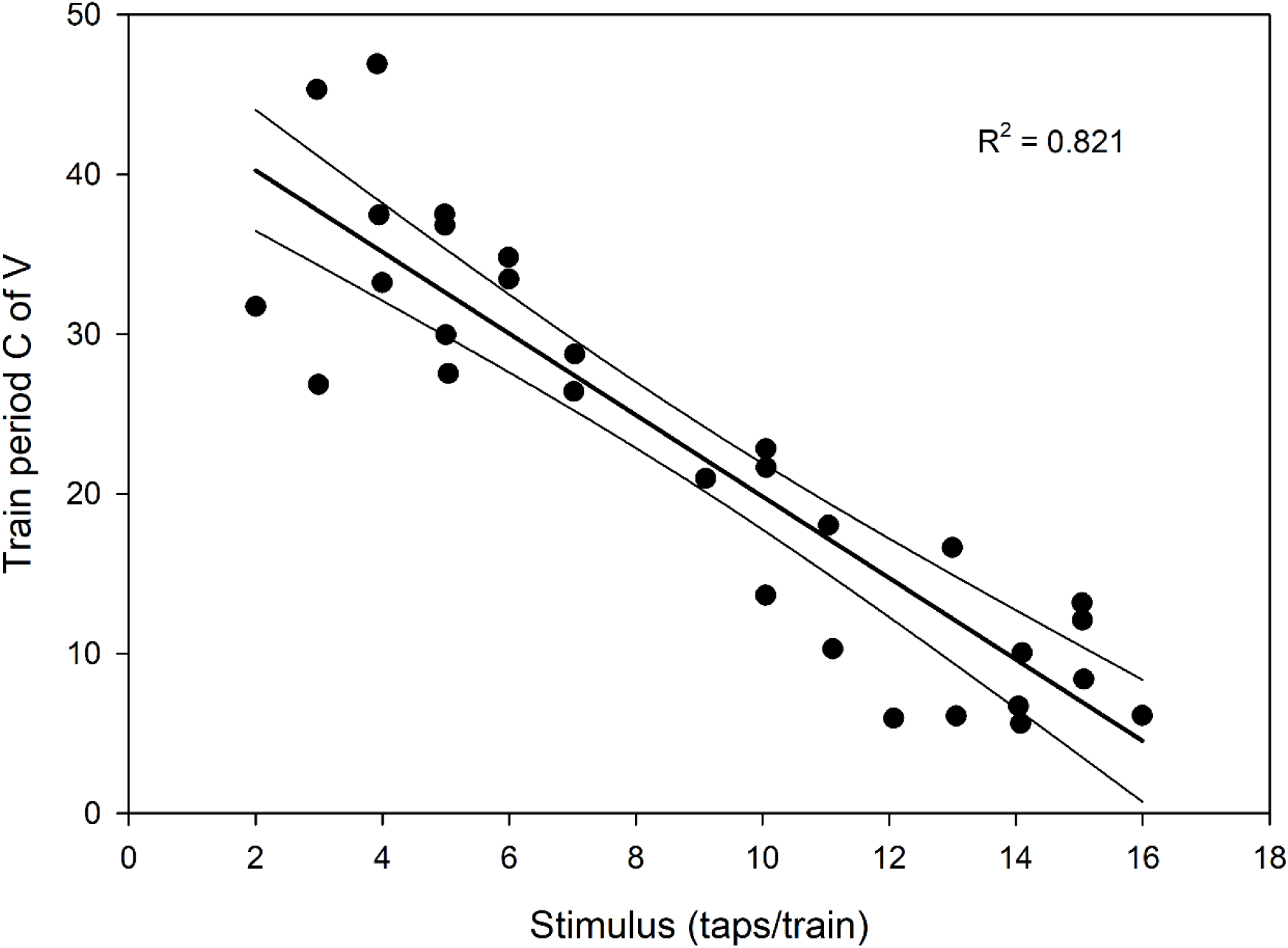
Relation between stimulus taps per train and stimulated male inter-train pause coefficient of variation. The increase in variation of inter-train pauses as stimulus tap-numbers decrease (i.e. approach the typical tap-number of the female signal) reflects increasing phonatactic behavior by the males. See text. n = 30 sessions, r^2^ = 0.82, F_1,28_ = 128.72, *P* < 10^-6^; Y = 45.36-2.553*X.

Consequently, as shown in Table 3, the data for male tapping parameters were split into responses to stimuli characteristic of females (< 8 taps/train) and those unambiguously characteristic of males (> 9 taps/train). Fig. 5 shows the probability distributions for all combinations (female, stimulated; male, unstimulated; male, stimulated at < 8 or > 8 taps/train; and male in the poststimulus state, after being stimulated at < 8 or > 8 taps/train).

**Figure 5.**
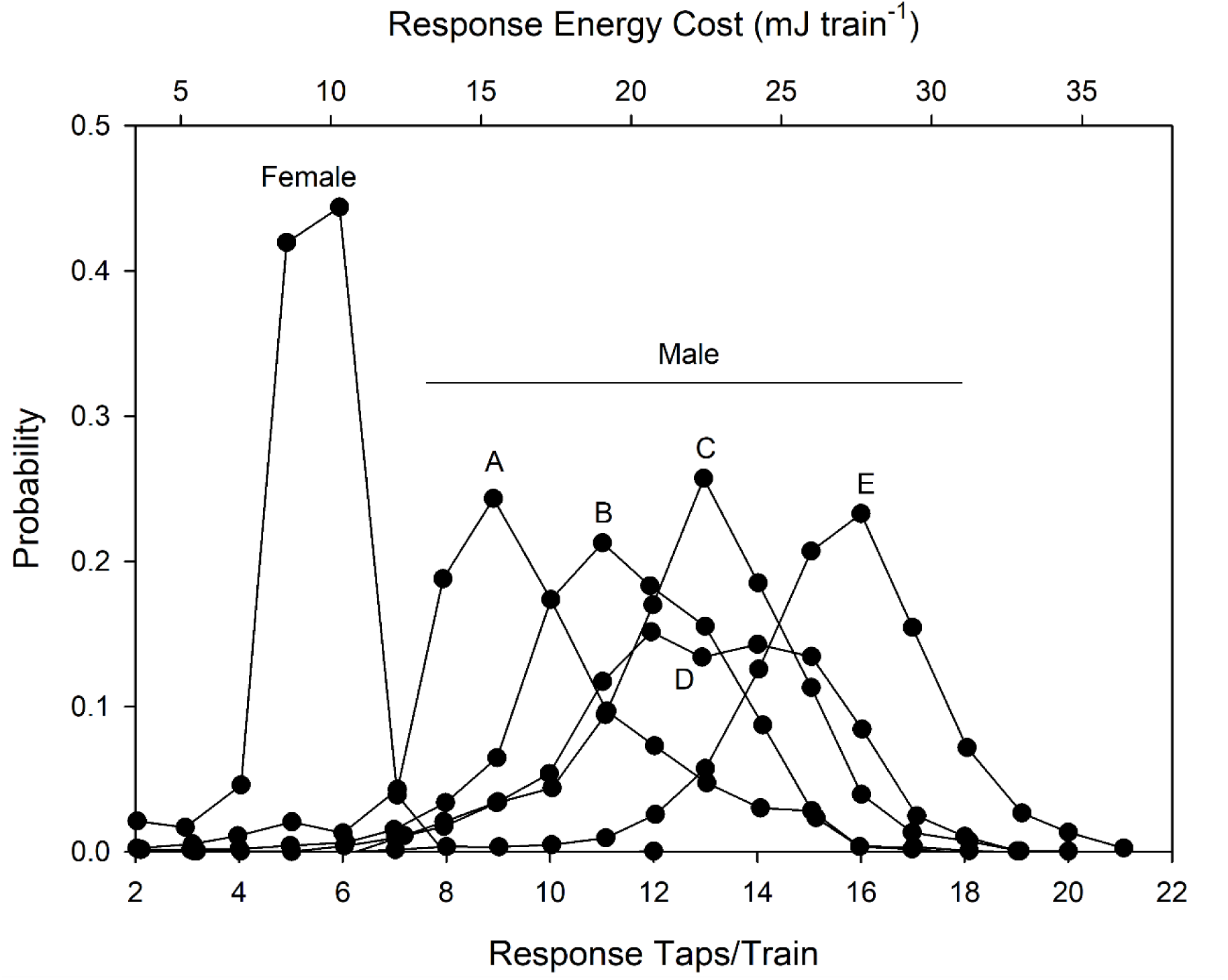
Combined tap-number probability distributions. Female distribution is labeled; they did not tap unless stimulated. Other distributions are male. A = unstimulated. B = poststimulated [S<8, where S = number of stimulus taps]. C = post-stimulated [S>9]. D = stimulated [S<8, characteristic of female tap number], the broad distribution reflects phonotactic behavior; see Fig. 3. E = stimulated [S>9, characteristic of male tap number]. See text.

**Figure 6.**
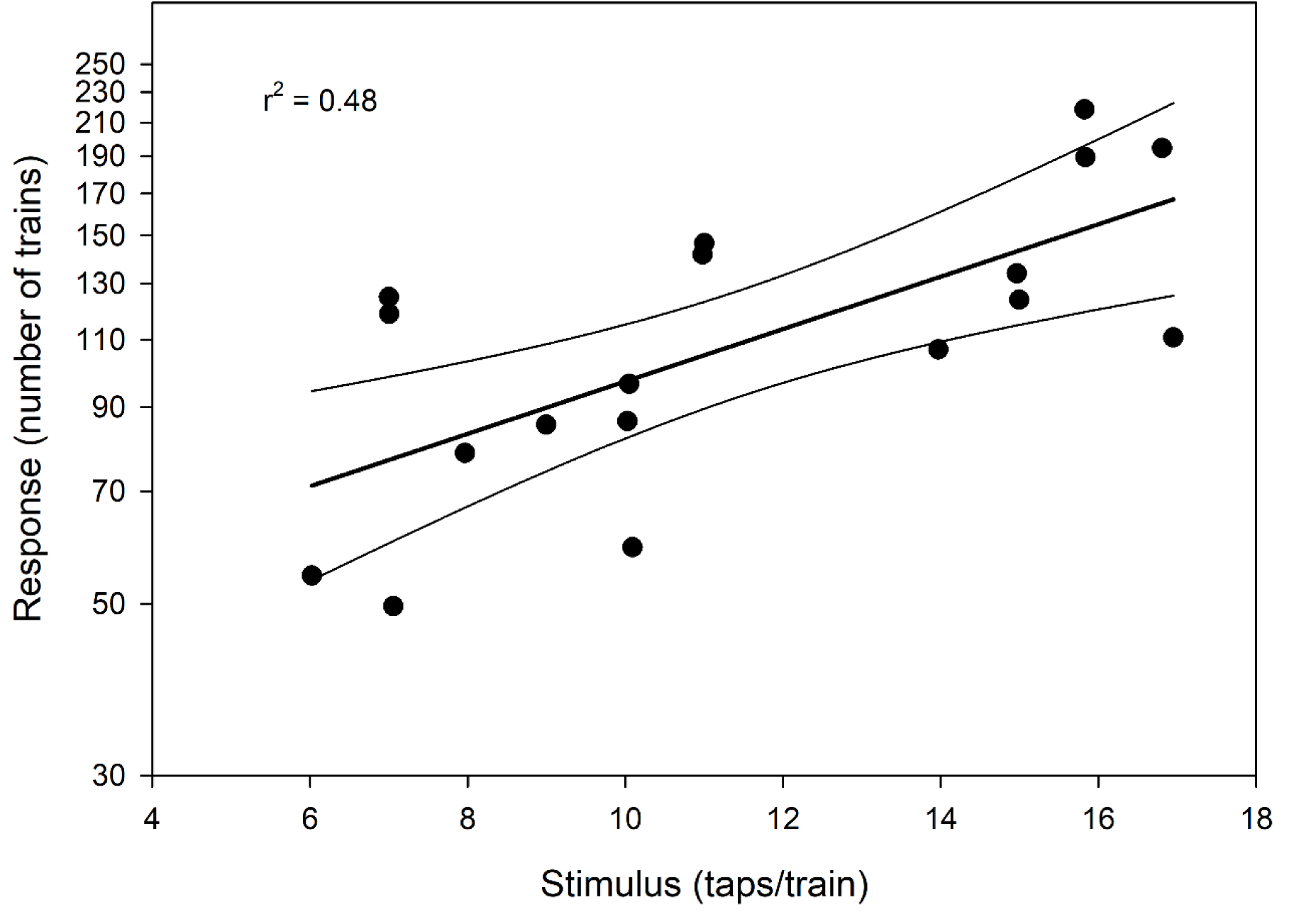
Exponential relation between stimulus tap-number and number of female response tap-trains while stimulus is applied. n = 18 sessions, r^2^ = 0.48, F_1,16_ = 14.65, *P* = 0.001; Y = 44.5*X^0.34^.

The intensity of female response to a tapping stimulus, as assayed by the number of total response tap-trains per monitoring period, was a strong, indeed exponential, function of the number of taps per stimulus train. Females did not respond at all to stimulus tap-trains containing fewer than 6 taps, and responded with six-fold greater intensity to tap-trains near the top of the male range (N = 16 - 18 taps/train) than to those near the bottom of the male range (6 - 8 taps/train). Fig. 6 shows the pattern.

In addition, females replied to stimulus tap-trains significantly more rapidly as the numbers of stimulus taps/train increased (Fig. 7). The pause before responding to a stimulus was always much longer for males than for females and varied according to stimulus conditions, as shown in Fig. 8.

**Figure 7.**
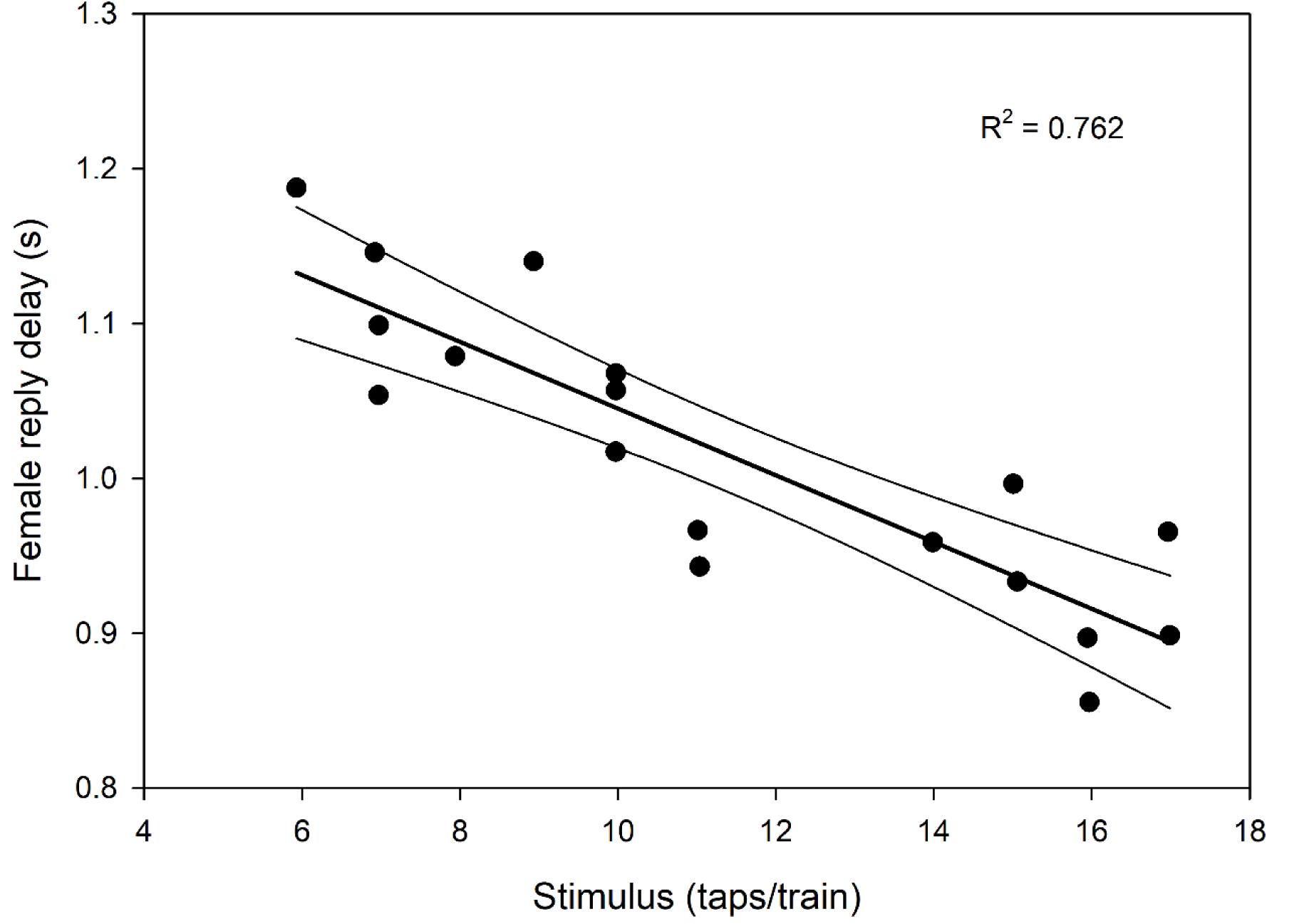
Effect of stimulus tap-number on female pause before replying to a stimulus (s). n = 18 sessions, r^2^ = 0.76, F_1,16_ = 51.13, *P* < 10^-6^; Y = 1.261-0.02157*X. See text.

**Figure 8.**
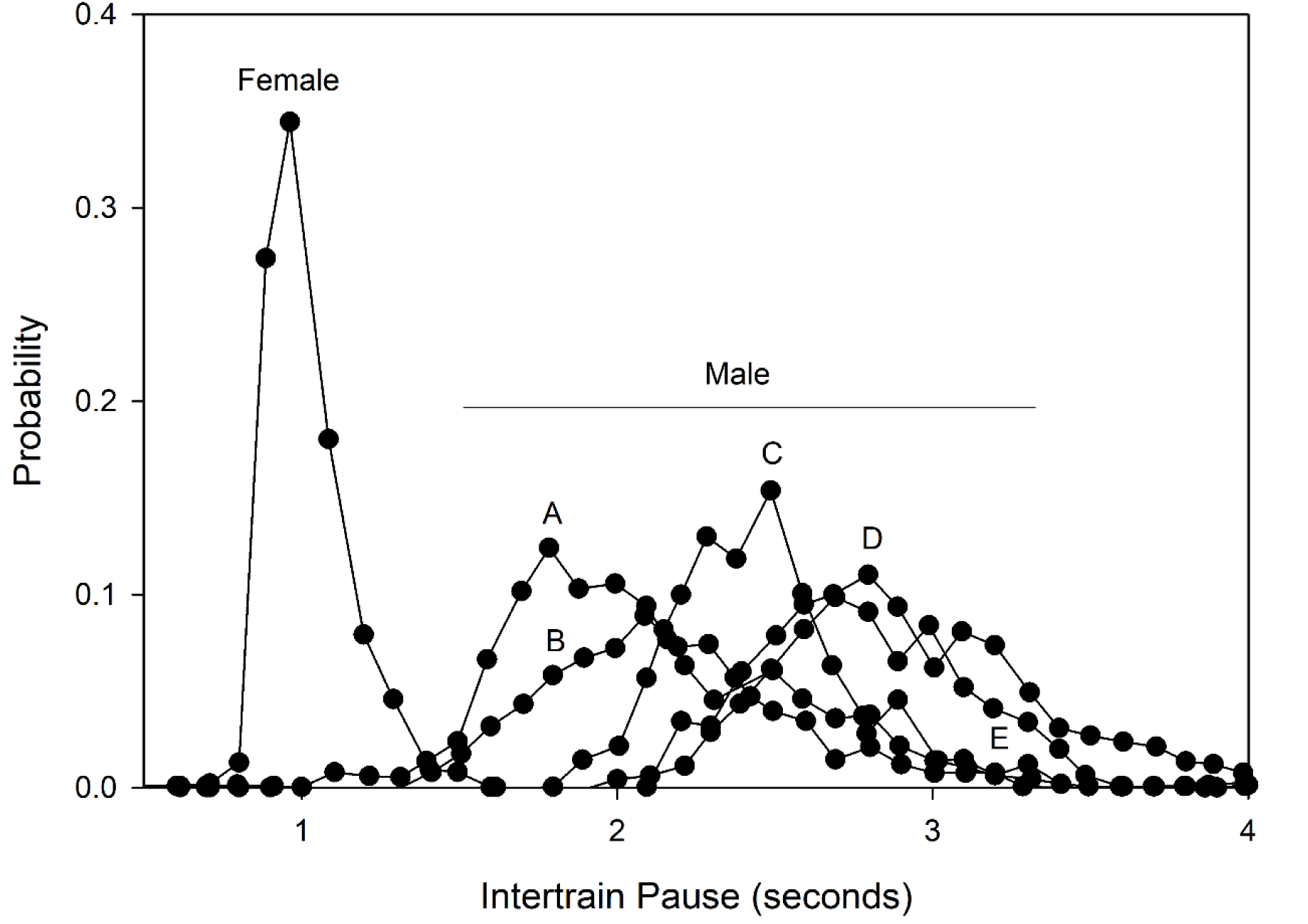
Combined probability distribution of male and female inter-train pauses. Female distribution is labeled. Others are male. A = stimulated [S>9]. B = stimulated [S<8]. C = unstimulated (note intermediate position). D = post-stimulated [S<8]. E = post-stimulated [S>9].

In contrast to the poststimulated males (see below), the females tapped a total of only 27 ± 22 times during the poststimulus period. Female poststimulus data have therefore been ignored in this analysis, as they were unlikely to be statistically meaningful. However, they do suggest that the females, though they do not spontaneously initiate tapping, may attempt to reestablish contact with another beetle with which they were communicating and which has recently stopped tapping.

### “Natural” male-female tapping behavior

Two male-female duets which took place in a partitioned monitoring tank, allowing only acoustic contact between the beetles, were monitored. These interchanges, which lasted 11 and 18 minutes, consisted respectively of 1651 and 2973 taps (224 and 372 trains). Data from these interchanges were combined, as no significant differences in taps/train, response time, *etc.* were found between the two. Plotting a small portion of one interchange as taps per train with a time-directed X axis reveals the essential distinctions between male and female signals quite well (Fig. 9). In particular, note how the female trains contain fewer taps and follow very closely on the male signals.

**Figure 9.**
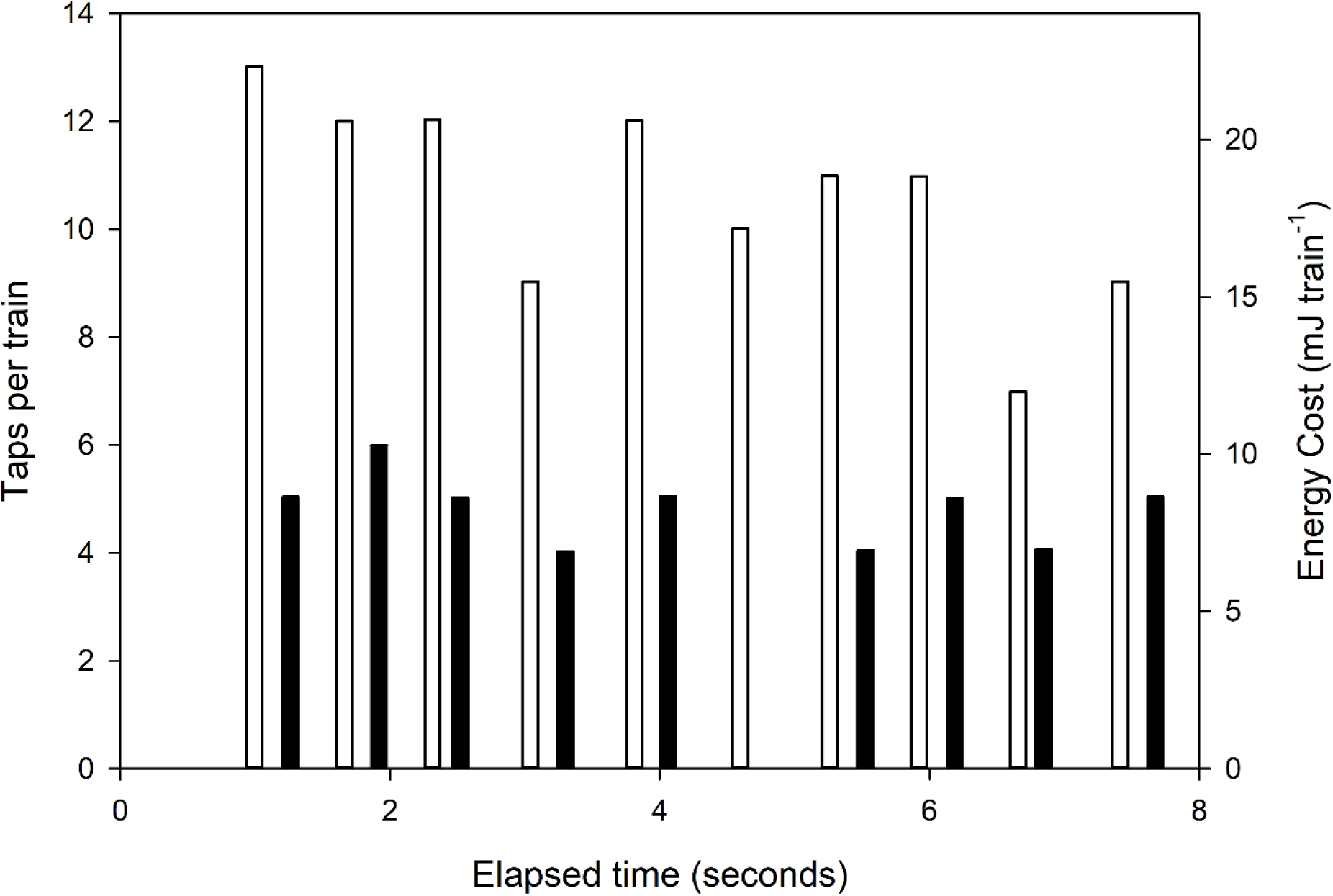
Representation of a small portion - 19 trains - of a male-female ‘conversation’. The brief female replies (filled bars) follow shortly after the longer male calls (empty bars). Note that the widths of the bars are not proportional to the durations of the tap-trains.

## DISCUSSION

### Male-female communication

In *P. striatus*, the female operates as a transponder rather than as an initiator. In any dual signaling system, biological or not, the ultimate object of which is direction-finding and co-location, it is logical for one partner to stay in place and respond to “ping” signals, while the other dynamically initiates those responses and then employs them to locate the transponder. Consequently, the transponder replies quickly while the initiator waits until the probability of a reply has declined to near-zero before sending another “ping.” This is seen clearly in the male post-stimulus distribution in Fig. 5, and probably explains the lack of spontaneous tapping by females. For the female to act as a transponder, responding only when stimulated and with a distinctive, low-redundancy, low-energy, minimal-delay signal to the high tap-numbers characteristic of male tapping, may allow her to allocate more of her energy budget for essentials such as producing viable eggs. This may carry a selective advantage in a xeric environment where energy must be conserved, whether because of energy or water limitation (Louw and Seely, 1982; Edney, 1971). The narrowly defined nature of the female’s response may also be important in species recognition. Such precise timing is a property of this form of communication which is predictable on theoretical grounds (the spectral information content of a tap, as opposed to an airborne signal such as a stridulation, is negligible). Thus, male beetles produce a variable, broad-band signal which is significantly affected by the presence, absence, nature and even the recent history of suitable acoustic stimuli, while the female responds with an unambiguous transponder/locator signal - if she responds at all.

### Male-male communication

The tap-happy male behavior seems intuitively counteradaptive. On energetic grounds alone, profligate communication behavior should be selected against. Indeed, when the beetles are kept in captivity, tapping among males is an almost round-the-clock activity. These interactions are not dissimilar to the “rival”s songs’ discussed in Dumortier (1963), which, however, are aggressive and territorial. No aggressive encounters were seen to take place between males in this species, though when two tapping males meet, one will usually try to mount the other. This behavior is identical to that used by males when mounting females and is likely triggered by the size and shape of the other beetle, as this species shows negligible sexual dimorphism and, judging from its tiny eyes and small number of ommatidia, its vision appears to be rudimentary. Observation suggested that these beetles are only aware of each other at very close range (2-4 cm) unless tapping was involved (author’s unpublished observations).

The phenomenon of specific interactions between males in communicating insects has received attention (see especially Spooner 1968; Alexander 1968, 1975; Lloyd 1979; Slobodchikoff and Spangler 1979; Hill 2009), but not, to my knowledge, in situations similar to these, where the call contains sex-coding data. Here, energy is expended in communication with a beetle which not only cannot directly transfer the beetle’s haploid complement to the next generation, but is likely to act as a competitor in such a transfer. Nevertheless, it may have a rationale.

It is obvious that the fixed, distinct nature of female trains facilitates the mutual recognition of males. Specifically, communication between males serves to increase the number of taps in each train, leading to a still greater differentiation between male and female calls, ensuring that a female signal will be immediately recognized if it occurs during inter-male communication.

Males are thus operationally aware of the sex of their communication partner, as evidenced by their phonotactic behavior, which is exclusively induced by calls characteristic of female beetles (< 8 stimulus taps per train; see Fig. 4). This is energetically expedient. Engaging in phonotactic behavior to locate other males would be genetically unproductive and wasteful of energy. In addition, inter-male phonotaxis would lead to aggregation of the males, reducing the area covered by their signals, and thus - assuming a random distribution of females - reducing the probability that an individual male would attract a reply from a passing female. Such a phenomenon may in fact occur in the tenebrionid beetle *Eupsophulus castaneus* (Slobodchikoff and Spangler 1979). It is perhaps significant that in that species, the females do not tap (ibid). Thus *Eupsophulus* females cannot be phonotactically located by the males, which aggregate when calling.

If the tap-number probability distributions are used to calculate the probability that a signal from one distribution is a member of another, this effect is clearly seen. Thus the probability of a signal characteristic of a female call occurring in an unstimulated male call can be estimated at 0.1058, whereas the corresponding probability for an inter-male call is only 0.0289 - in other words, inter-male calling increases the sex-coding of a tapping signal by almost 4-fold, even if one ignores the two-fold difference in speed of response to stimuli (1.03 second for females *vs.* >= 2.09 seconds for males) which presumably adds some information to the process of sex recognition. In a sense, the males’ behavior is more to the females’ than the males’ advantage if the female does indeed act on this information.

These phenomena lead one to hypothesize that the female will respond preferentially to large tap per train numbers, as they in turn imply not only greater fitness but the likely presence of more than one male, with a correspondingly increased probability of fertilization and, perhaps more importantly, of inter-male competition. The number of female response trains does indeed increase exponentially with increasing numbers of taps per stimulus train (Fig. 6). One might infer that such energetic signaling would handicap the male beetle by making him more conspicuous to potential predators. If so, the selective advantage of elevated female responsiveness apparently outweighs that handicap (for a general discussion, see Grafen 1990 and references therein).

### The “communication circle”: a speculative model

Male-male interactions may have another effect. A tapping beetle transfers acoustic energy to the ground, and this energy radiates omnidirectionally from its source, creating what one might call a ‘communication circle’. The acoustic energy relevant to the beetles’ communication travels along the surface of the ground in the form of Rayleigh waves to which tarsal receptors in arthropods are known to be sensitive (Autrum 1963). Absolute communication distances in these beetles are uncertain; let us call the radius of the circle of communication r, and set its value equal to unity. The area of the communication circle is then π radius units.

Obviously, two beetles cannot communicate if they are more than one radius unit apart. For communication to take place, their circles must interpenetrate and the total area their combined signals will cover must be smaller, and with it the probability of a female occurring in that combined area (though this may be partly offset if the distribution of these beetles is “patchy”, leading to an increased probability that a female will be present if two males are already present). If males are closer together, the probability that communication between them will occur begins to rise. At the same time, their shared area increases relative to the single-male state. It can be shown that

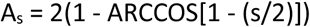

where A_s_ is the shared area, and s the beetle separation, in radius units. The proportional shared area reaches 0.33 when the beetles can potentially communicate, and rises to 1.00 when the two beetles are a negligible distance apart. It is immediately obvious that males communicating at the limits of their ranges have areas of “exclusive communication” equal to 0.67 of their circles of communication, in which they can receive replies from a female beetle without the other male’s knowledge. Moreover, the female will be stimulated by tap-numbers characteristic of inter-male communication, considerably increasing her responsiveness (Fig. 6).

It is therefore not to the males’ advantage to locate each other or otherwise attempt to aggregate once inter-male communication has started. Though this means that a male can participate if another male begins to communicate with a female that they can both hear, it also means that he may use a considerable amount of energy in the process of “homing”, and that he may encounter competition from the male closer to the female. Consequently the standardized shared area of the circles can be regarded as an index of potential competition, if the simplifying assumption is made that all interactions are between beetles with identical radii of influence. The potential competition index is zero if the males are physically far removed from each other (> 2 radii). However, its value becomes non-zero between 2 and 1 radii, *even before the beetles can became directly aware of each other*. This seeming paradox means that both beetles may now be within the signaling range of a third beetle replying to one or both of them, and are therefore potentially able to act as competitors even though they are not aware of each other. The value of this index reaches 0.33 when the beetles can become aware of each other directly, and climbs to unity when the beetles are a negligible distance apart and neither can communicate with a third beetle without the other’s knowledge.

At two or more radii apart, individual males have a consistent advantage owing to the absence of competition and increased area of exclusive communication, but the advantage is low owing to a) the relative infrequency with which they tap when not stimulated, and b) the female’s low responsiveness to the small tap-numbers characteristic of unsolicited male tapping. Then, as one intuitively expects, individual male advantage is sharply reduced as the males draw together. Any interchange initiated by one male with a female may be heard by the other male (for example, the female may be between them), while the effective tap-number remains in its unstimulated condition. The situation changes abruptly when the males can communicate. As soon as they start to do so, their tapping density rises from *ca.* 8 taps per minute to *ca.* 100 and their tap-number from *∼*9 to *∼*15, significantly increasing the probability of female replies. Even corrected for the distance-dependent probability of this interchange starting, the advantage now swings abruptly in the direction of the individual male. Thus, the individual male - thanks to inter-male communication - now has not only a much greater absolute chance of eliciting a response from a female, but also a distinct probability that this response will be imperceptible to the other male, and hence to his benefit alone. It could thus be argued that individual selection is a strong component in the maintenance of this behavior by natural selection. Even when competition is almost certain to occur (i.e. close to the other male), the mean benefit gained by inter-male tapping can in individual terms be reduced by only 50% if either male has an equal chance of displacing the other and mating with the female. This reduction is still a huge improvement over the unstimulated condition, and is therefore still to the advantage of the individual.

### The advantage to females of male-male communication

If the situation is now viewed from the female’s perspective, it is to her advantage to preferentially reply in the presence of more than one male. This is particularly true if the probability of inter-male competition is high (which can be increased if her sensitivity to signals is equal to that of the males’, but her signal can be heard over a larger radius, perhaps as a result of slightly heavier mean body mass in females [author’s unpublished observations]). This results in a correspondingly greater probability that the more robust and determined male will mate with her (see especially Reinhold et al. 1998). This may contribute little if anything to her short-term advantage, but will help to ensure that her genetic complement is paired to that of a beetle that has demonstrated a competitive edge over another, and therefore maximizes the probability that her offspring will successfully transfer her genetic complement to the next generation. In ultimate terms this is probably the driving force behind inter-male communication. Paradoxically the advantage gained by individual males, though perhaps significant, has only been gained within the framework of the female’s pattern of response - which may work to her benefit only if inter-male competition is an important factor in inter-male communication.

### Energy savings

The metabolic energy cost per tap is 1.725 mJ for a beetle weighing ∼3 g (Lighton 1987). The position of the tap in the tap-train has no significant effect on its cost as determined by kinematic analysis, which also revealed an efficiency, or power output as a percentage of metabolic input, of 23%, a typical value for muscle efficiency across a broad range of taxa (Mogensen et al. 2006 and references therein). For male beetles with a mean unstimulated tap-train of 9.63 taps train^-1^ and a mean number of tap-trains per group of 4.74 trains group^-1^ (see Table 1), sending a tapping message thus costs 78.7 mJ. If we assume a modest tapping communication range of 0.5 m and a circular reception area for substrate-borne vibrations, the 78.7 mJ energetic investment allows an area of 0.785 m^2^ to be searched for other beetles. This is equivalent to a search cost of ∼100 mJ m^-2^. Pedestrian locomotion, in contrast, can be reasonably assumed to allow direct detection of beetles no more than ∼5 cm to either side, creating a “swathe of contact” ∼10 cm wide. The minimum cost of transport in this species is known (∼100 mJ m^-1^ for a 3 g beetle; calculated from Lighton 1985). Each meter walked thus covers a search area of 0.1 m^2^, yielding a search cost of 100 / 0.1 or 1000 mJ m^-2^. From this, we estimate that the energy required for a beetle to search an area for a prospective mate via tapping communication is approximately 12.7-fold less than encountering another beetle via the *flaneur* method (walking). This does not consider the ∼50% probability that a beetle thus encountered is of the same sex, or the possibility that a female beetle is unreceptive. It is, however, complicated by the existence of male-male interactions that will add energy expenditure – from which it is reasonable if marginally Panglossian (Gould and Lewontin, 1979) to infer that such interactions confer benefits that outweigh their cost, as discussed above (“communication circle”).

### Future prospects

Several other tenebrionid beetles in the Molurini tribe such as *Moluris nitida* tap in ways similar to *Psammodes striatus*, though at different frequencies; others dynamically modulate their inter-tap durations and engage in long tap-trains that start at ∼1 Hz and reach a crescendo of >10 Hz (author’s unpublished observations). Quantifying these diverse substrate-borne communication protocols could lead to interesting insights, particularly if energy allocations are analyzed in conjunction with ecology and behavior.

## ACKNOWLEDGEMENTS

I thank the late Gideon Louw for the use of his laboratory at the University of Cape Town and for his support in many ways, great and small; the South African CSIR for financial support; Pablo Schilman for assisting with the figures; the novelist and linguist Menan du Plessis for her practical insights regarding the theory and practice of communication; the late George Bartholomew (“Bart”) for accepting me as his last graduate student at UCLA, where I gained access to state of the art metabolic measurement equipment; Peggy Hill, Karl Kaiyala and Robbin Turner for helpful comments on an earlier draft of the MS; and Sable Systems International for financial support during the writing of this paper via their Basic Science Initiative.

